# Identifying genetic regulations on immune cell type proportions and their impacts on autoimmune diseases

**DOI:** 10.64898/2026.02.26.708418

**Authors:** Chen Lin, Jiangnan Shen, Jiaxuan Sun, Yuhan Xie, Leqi Xu, Yingxin Lin, Jiaqi Hu, Hongyu Zhao

**Affiliations:** Department of Biostatistics, Yale School of Public Health, New Haven, CT, USA 06510; Department of Chronic Disease Epidemiology, Yale School of Public Health, New Haven, CT, USA 06510

## Abstract

Genetic regulation of immune cell composition plays a crucial role in the etiology of complex diseases, yet remains poorly understood. We propose a unified analytical framework that integrates genome-wide association studies (GWAS) of cell type proportions with cell-type-wide association studies (cWAS) to systematically characterize both the genetic regulation of immune cell composition and its downstream effects on disease risk. Using single-cell RNA sequencing data from the OneK1K cohort, we conducted a GWAS of immune cell-type proportions with a depth-weighted quasi-binomial model designed for bounded, overdispersed traits. We identified 47 genome-wide significant loci influencing eight fine-labeled immune cell subtypes. Leveraging these identified genetic effects, we further imputed genetically regulated proportions (GRPs) using polygenic risk score (PRS)-based imputation and assessed their associations with complex diseases through cWAS. We identified five significant cell type-disease associations, including two with type 1 diabetes, two with Crohn’s disease, and one with ulcerative colitis. Together, our results demonstrate that cell type proportions observed in scRNA-seq can reveal regulatory loci and offer insights into how genetic variations regulate immune cell type proportions to affect disease risk. Although we focused on immune single-cell data, our framework is applicable to other tissues or cellular compositions as scRNA-seq datasets expand.

**Author Summary:** Genome-wide association studies (GWASs) have uncovered many disease-associated signals, yet most lie in noncoding regions and are difficult to interpret. Mapping GWAS signals to the relevant cell types is therefore important for better understanding the biological mechanisms that drive disease. A major challenge is that observed gene expression and measured cell-type proportions can be influenced by environmental factors and disease status. In contrast, genotypes are less affected by these factors, making them more reliable for interpreting factors of diseases. Moreover, the cell-type proportions are bounded and often skewed, so standard GWAS models that rely on Gaussian assumptions may lose power. To address this, we developed a quasi-binomial approach that better matches the data and improves discovery while controlling false positives. In real data, our method identified more genetic loci associated with cell-type proportions than a traditional linear model. To further investigate how genetic variation regulates immune cell composition to influence disease risk, we integrated our results with disease GWAS summary statistics to identify immune cell types that may contribute to disease susceptibility. Together, our results link disease-associated GWAS signals to specific immune cell types and provide insights into the cellular mechanisms that may underlie these diseases.

## Introduction

Genome-wide association studies (GWASs) have identified thousands of genetic variants associated with complex diseases. However, most of these variants are in non-coding regions, making it challenging to interpret their biological functions. Previous studies show that genetic regulation of complex traits is closely linked to specific tissues and cell types^1,2,3^. Consequently, characterizing genetically regulated variation in cellular composition may enable the identification of trait-relevant cell types and the revelation of underlying biological mechanisms.

Many studies have attempted to link GWAS signals to relevant cell types by integrating transcriptomic data^4,5,6^. However, most of these efforts rely on observed gene expression levels or cell-type proportions, both of which can be influenced by environmental factors and disease status. This introduces confounding and potential reverse causation, which can bias association estimates. In contrast, genotypes are less influenced by environmental factors and disease status, making them more reliable for interpreting disease-associated factors. Cell-type-wide association study (cWAS) was developed to estimate the contribution of genetically regulated cell type proportions (GRP) to human diseases^7^. While cWAS has advanced our understanding of disease-cell type associations, it relies on deconvolution of bulk RNA-seq data and cannot identify genetic variants that directly regulate the measured cell type proportions.

The emergence of large-scale single-cell RNA sequencing (scRNA-seq) datasets now enables direct identification of genetic effects on cell-type proportions across individuals and provides a robust approach to investigate the associations between GRP and complex diseases. For example, a recent GWAS was conducted on proportions of immune cell types using scRNA-seq data, yielding only a limited number of significant signals^8^. One key limitation is that cell-type proportions derived from scRNA-seq data are bounded, often bulk-skewed, and overdispersed due to biological and technical variability. These features violate the assumptions underlying traditional linear GWAS models, thereby reducing power.

In this study, we introduce a framework to investigate the genetic regulation of cell-type proportions and their associations with complex diseases. The key advance is to treat scRNA-seq–derived cell-type proportions as bounded, overdispersed traits measured with depth-dependent precision, rather than assuming a Gaussian distribution. We propose a quasi-binomial regression model that improves power while remaining well-calibrated for false-positive controls. Applied to the OneK1K^9^ dataset of peripheral blood mononuclear cells (PBMCs), the quasi-binomial model identified 12 more genome-wide significant loci associated with cell-type proportions than the standard linear model. Based on the derived cell-type proportion GWAS summary statistics, we imputed GRPs using the polygenic risk score (PRS) method and tested for association with disease phenotypes via cWAS. Our analysis uncovered novel associations between GRPs and immune-related diseases, offering insights into potential causal cell types and mechanisms of these diseases. The framework connects GWAS signals for disease to immune cell types and provides insight into how genetic variants modulate immune cell composition to influence disease susceptibility.

## Results

### GWAS analysis of cell type proportion on the OneK1K dataset

To identify the genetic regulation of cell-type proportions, we applied our proposed depth-weighted quasi-binomial model on the OneK1K dataset, which includes scRNA-seq data from 1.27 million PBMCs obtained from 981 healthy human donors of European ancestry, including 416 males and 565 females^9^ (Methods). Our study focused on 19 fine-labeled cell types (Table S1; Fig S1), and further integrated them into six broader cell types, including B cells, CD4 T cells, CD8 T cells, Other T cells, Monocytes, and Natural Killer cells (Table S2; Fig S1). Seventeen donors with abnormally large cell type proportions were removed from the study as outliers (Methods).

We identified 47 genome-wide significant genetic variants (*p* < 5 × 10^−8^) associated with at least one of eight cell subtypes (B intermediate, CD14^+^ monocytes, CD16^+^ monocytes, CD4CTL, CD4TEM, NKCD56bright, Plasmablast, and MAIT), with 12 independent lead SNPs after filtering out those in high linkage disequilibrium (Table 1; Table S3). However, no significant associations were detected when analyzing coarser cell categories (Fig S3), likely due to the loss of granularity in cellular heterogeneity and cell-type-specific genetic effects. In contrast, the linear model only identified 35 significant genetic signals associated with at least one of six distinct cell types (B intermediate, CD14^+^ monocytes, CD16^+^ monocytes, CD4CTL, NKCD56bright and Plasmablast), with 8 independent lead SNPs after LD clumping (Table 1; Table S3). Notably, all 35 variants detected by the linear model were fully encompassed by those detected by the quasi-binomial model, highlighting the quasi-binomial model’s higher power and robustness for detecting genetic associations with cell type proportions.

**Table 1.** Lead SNPs from quasi-binomial model results.

| ct_type | CHR | BP | Effect Allele | Quasi-binomial model p-value | Linear model p-value |
| --- | --- | --- | --- | --- | --- |
| Bintermediate | 14 | 92826246 | A | 5.288e-09 | 3.668e-08 |
| CD14<sup>+</sup> monocytes | 10 | 3094356 | G | 1.958e-08 | 4.335e-08 |
| CD16<sup>+</sup> monocytes | 2 | 177494043 | C | 2.514e-08 | 3.085e-08 |
| CD16<sup>+</sup> monocytes | 10 | 126178267 | T | 7.249e-09 | 6.026e-08 |
| CD16<sup>+</sup> monocytes | 11 | 75288152 | G | 8.939e-10 | 3.043e-09 |
| CD4CTL | 7 | 138798087 | A | 3.487e-11 | 5.929e-10 |
| CD4CTL | 21 | 16270327 | T | 1.536e-08 | 1.072e-07 |
| CD4TEM | 6 | 142621947 | G | 4.660e-08 | 5.188e-08 |
| NKCD56bright | 2 | 111846068 | T | 1.669e-08 | 2.678e-08 |
| NKCD56bright | 12 | 71853712 | T | 6.687e-10 | 1.353e-09 |
| Plasmablast | 6 | 43715425 | A | 4.370e-10 | 1.044e-09 |
| MAIT | 3 | 175257167 | C | 3.425e-08 | 2.257e-07 |

We further compared quasi-binomial and linear GWAS results across cell types. For the CD16^+^monocytes (CD16Mono), association evidence was highly concordant across SNPs between the two models (Pearson r = 0.998; Fig 2A), while a distinct subset demonstrated substantially stronger statistical significance under the quasi-binomial model, suggesting improved power for association detection. For example, variant rs4962627 reached genome-wide significance (*p* < 5 × 10^−8^) in the quasi-binomial analysis (*p* = 7.25 × 10^−9^) but not in the linear model (*p* = 6.03 × 10^−8^). For the NK CD56 bright cells, quasi-binomial regression identified three additional genome-wide significant variants relative to the linear model (Fig 2B). Furthermore, QQ plots for the quasi-binomial and linear models showed no evidence of inflation (Fig S4∼S5), suggesting that both methods-controlled type I error appropriately. These results underscore the advantages of quasi-binomial regression for identifying genetic effects on cell-type proportions.

**Fig 1.**
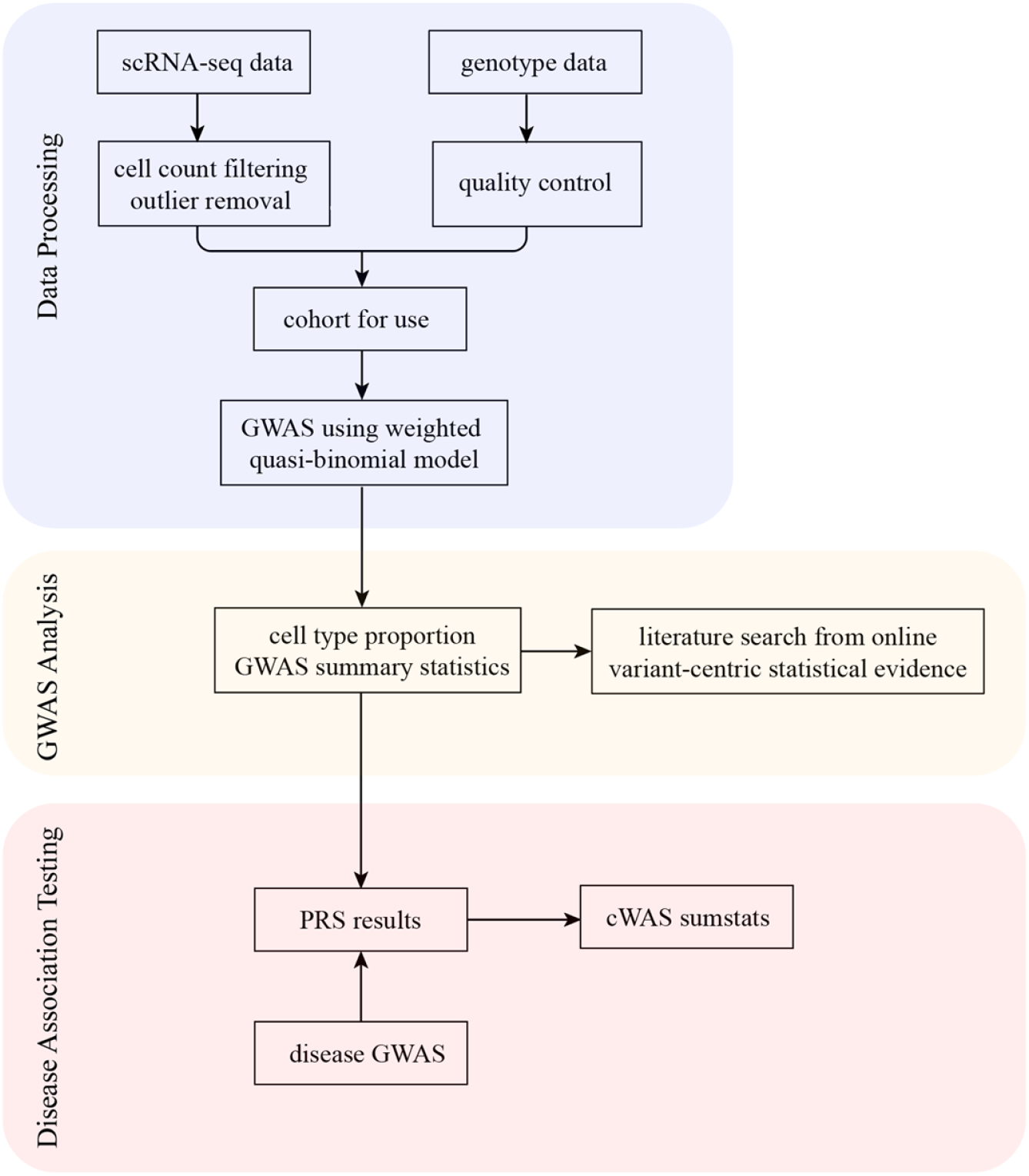
Overview of the study design for identifying genetic regulation of immune cell type proportions and their disease associations. **a**, Data Processing. Single-cell RNA-seq (scRNA-seq) data were processed to filter out low-quality genes, cells with extreme cell type proportions, and technical outliers. Individuals with sufficient sequencing depth were retained for genome-wide association studies (GWAS). **b**, GWAS Analysis. A weighted quasi-binomial model was applied to test associations between genetic variants and proportions of immune cell types. Binomial enrichment tests were then used to identify disease-relevant cell types. Literature-based evidence was gathered to support biological interpretations. **c**, Disease Association Testing. Polygenic risk score (PRS) models were trained using cell type GWAS results and applied to disease GWAS summary statistics. Cell-type Wide Association Studies (cWAS) were performed to identify significant genetically regulated cell type proportion (GRP) associations with autoimmune diseases.

**Fig 2.**
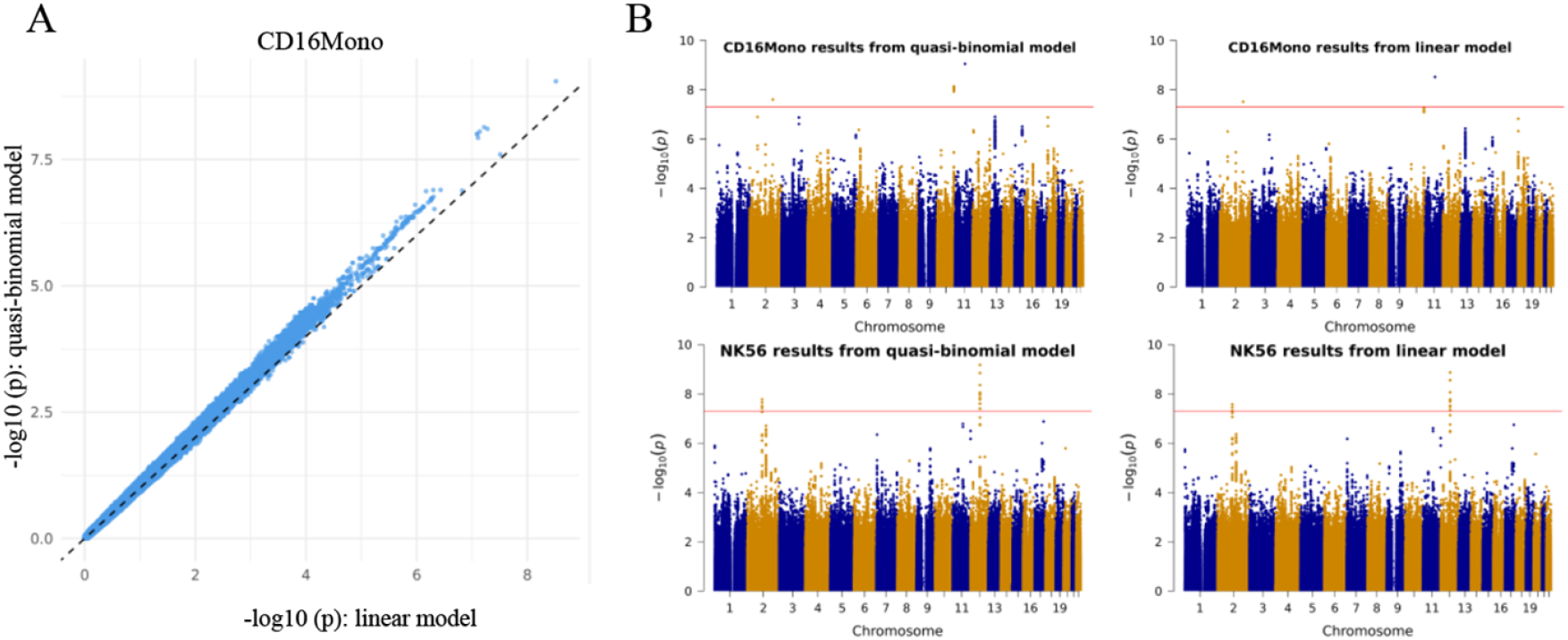
Quasi-binomial regression identified more GWAS signals for immune cell-type proportions. (A) Comparison of linear model and quasi-binomial model association p-values for CD16^+^monocytes. (B) Manhattan plots of CD16^+^monocytes and NK56 GWAS results using quasi-binomial model and linear model. The red horizontal line indicates the genome-wide significance threshold (p = 5 × 10^−8^).

### GWAS signals for cell type proportions are molecular and phenotypically relevant

We further assess the molecular and phenotypic evidence for the lead composition quantitative trait loci (QTLs) identified in the quasi-binomial model. Specifically, we searched variants of interest in the Open Targets^10^ platform, a database aggregating genetic association evidence from the GWAS Catalog, large-scale biobank GWAS summary statistics, and Phenome-wide association study (PheWAS) analyses, for functions of nearby genes, relationships with eQTLs, and pleiotropic associations.

The first example SNP, 2/27/2026, was significantly associated with the NK CD56bright cell proportions (Fig 3A) in both models (*p*_linear_ = 4.87 × 10^−8^; *p*_quasi_ = 3.12 × 10^−8^). It is proximal to *BCL2L11*, a critical regulator of apoptosis essential for immune cell homeostasis^11^, suggesting a potential role for apoptotic pathways in NK cell differentiation. PheWAS analysis further connects this variant to mean corpuscular hemoglobin levels, consistent with previous reports identifying GWAS lead variants in this genomic region associated with hematological traits^12,13^. Consequently, these findings indicate that rs56226558 modulates NK CD56bright cell proportions through regulatory mechanisms affecting cell survival, proliferation, or differentiation, with potential implications for immune function and hematopoiesis.

**Fig 3.**
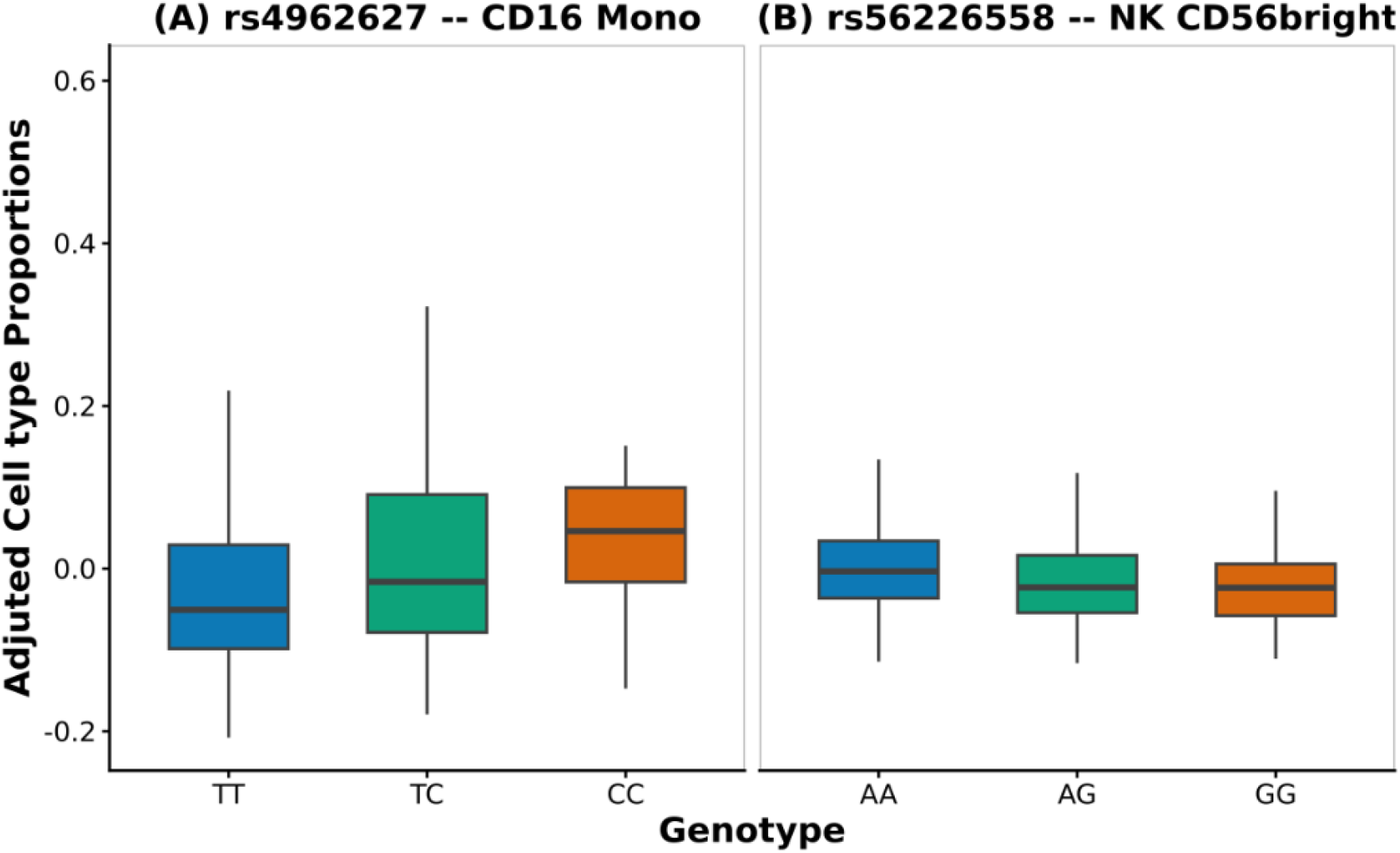
Genotype-dependent variation in adjusted cell-type proportions. Boxplots show residual CD16+ monocyte and NK CD56bright cell proportions after adjusting for covariates (age, sex, PCs), stratified by genotype at rs4962627 and rs56226558, respectively. These distributions illustrate genotype-specific regulatory effects on immune cell composition.

The second example SNP, rs4962627, located near *LHPP*, is associated with increased proportions of CD16^+^monocytes (Fig 3B) and reached genome-wide significance in the quasi-binomial analysis (*p*_quasi_ = 7.25 × 10^−9^) but not in the linear model (*p*_linear_ = 6.03 × 10^−8^). *LHPP* encodes a protein histidine phosphatase involved in phosphohistidine metabolism, a pathway implicated in tumor suppression, cellular signaling, and protein regulation^14^, and has been previously linked to white blood cell counts, neutrophil counts, and serum phosphate levels^15^. rs4962627 is also an eQTL for *EEF1AKMT2*, a gene involved in urate metabolism and phosphate regulation^15^, suggesting coordinated regulation of immune and metabolic traits. Given the pro-inflammatory and tissue-recruiting functions of CD16^+^monocytes, this locus may regulate phosphate metabolism in shaping monocyte differentiation. Additionally, PheWAS analysis suggests its association with the prevalence of *TestASV_11*^*16*^, a microbial taxon with the Lachnospiraceae family, implicated in immune regulation and inflammation, pointing to potential microbiome-mediated effects^17,18^. Together, these findings suggest that rs4962627 modulates CD16^+^monocyte abundance through integrated effects on metabolism, gene regulation, and host–microbiome interactions.

In conclusion, these examples reveal that many lead SNPs not only reside in proximity to genes with known immune or hematopoietic functions, but also act as eQTLs and show significant associations with relevant molecular traits and clinical phenotypes. These findings suggest that the genetic regulation of immune cell type proportions may involve transcriptional and post-transcriptional mechanisms, with broader implications for complex traits and diseases.

### Associations between GRPs and autoimmune diseases

To further study the associations between GRPs and diseases, we imputed GRPs from GWAS summary statistics and tested their associations with eight autoimmune diseases (Methods; Table 2). We identified five significant cell type-disease associations after controlling the false discovery rate using the Benjamini-Hochberg correction (FDR < 0.1, Fig 4; Fig S6). Among them, Crohn’s disease showed negative associations with CD16^+^monocytes and NK CD56bright cells, indicating that lower genetically predicted levels of these cell types were associated with higher Crohn’s disease risk (Fig 4).

**Table 2.** GWAS Summary Statistics of Autoimmune Diseases.

| Abbreviation | Disease | GWAS Sample Size |  |
| --- | --- | --- | --- |
|  |  | # of Cases | # of Controls |
| AS<sup>19</sup> | Ankylosing spondylitis | 495 | 371238 |
| SLE<sup>20</sup> | Systemic Lupus Erythematosus | 5201 | 9066 |
| RA<sup>21</sup> | Rheumatoid Arthritis | 2843 | 5540 |
| MS<sup>22</sup> | Multiple Sclerosis | 14802 | 26703 |
| IBD<sup>23</sup> | Inflammatory Bowel Disease | 25042 | 34915 |
| Crohn<sup>23</sup> | Crohn's Disease | 12194 | 28072 |
| UC<sup>23</sup> | Ulcerative Colitis | 12366 | 33609 |
| T1D<sup>24</sup> | Type 1 Diabetes | 9266 | 15574 |

**Fig 4.**
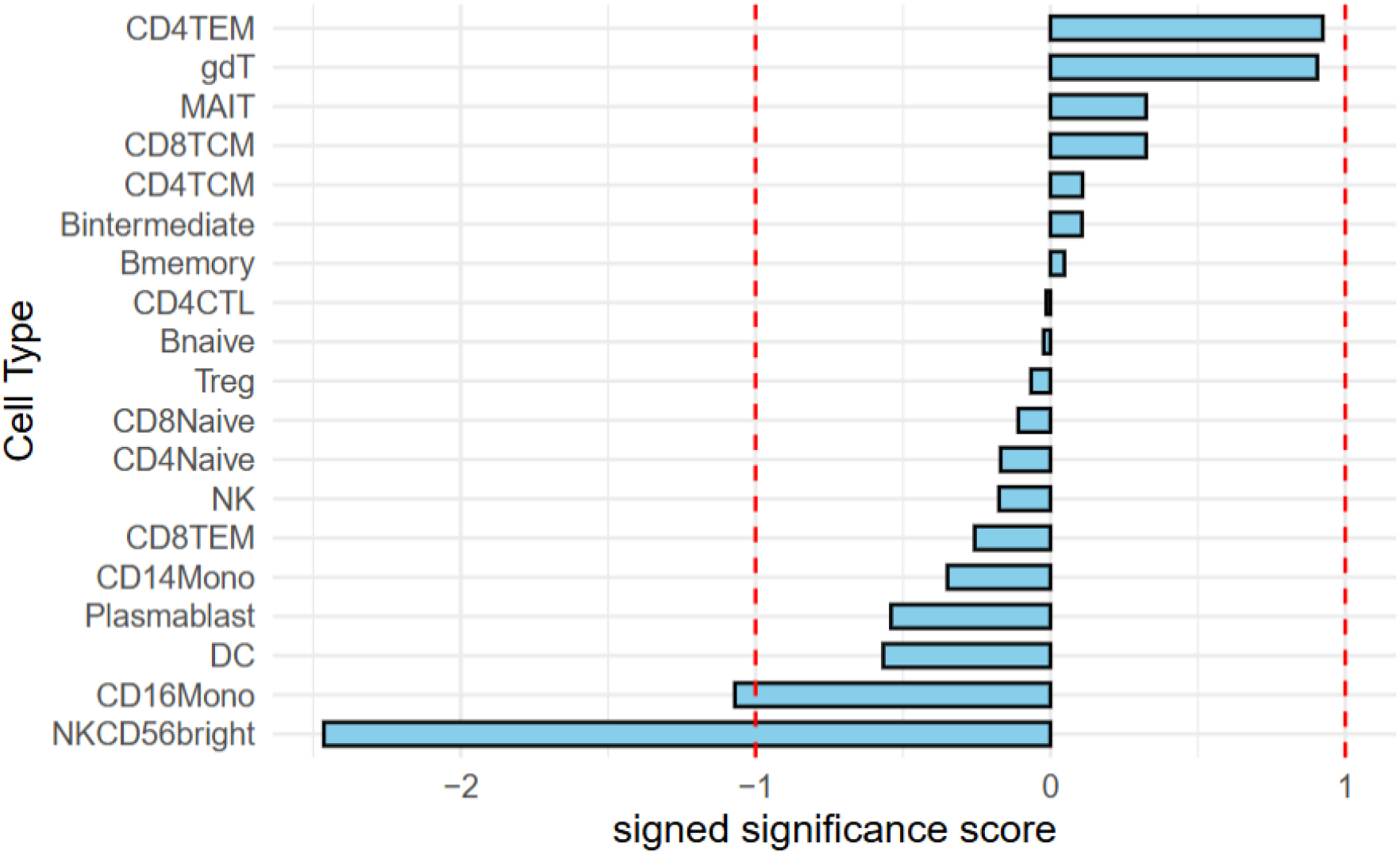
cWAS results for Crohn’s Disease in whole blood. The plot displays signed significance scores for the association between genetically regulated cell type proportions and Crohn’s Disease risk. The x-axis represents the signed significance score, defined as the negative log10 of the FDR-adjusted p-value multiplied by the sign of the corresponding Z-score. Positive values indicate increased proportions associated with elevated Crohn’s Disease risk; negative values indicate decreased proportions. Dashed red lines mark the FDR threshold of 0.1.

This finding aligns with previous research highlighting the role of CD16^+^monocytes and NK CD56bright cells in Crohn’s disease pathogenesis. For instance, CD16^+^monocytes contribute to the inflammatory infiltrate in Crohn’s disease mucosa and have been implicated as pro-inflammatory effector populations in inflammatory bowel disease^25^. The negative correlation with circulating CD16^+^monocytes is consistent with the possibility that genetically influenced differences in trafficking or tissue recruitment may shift these cells from blood to the inflamed gut, thereby linking systemic GRPs to mucosal immunopathology.

Moreover, NK CD56bright cells are primarily known for their immunoregulatory functions, secreting high levels of cytokines while exhibiting limited cytotoxicity^26^. Specifically, intestinal mucosal NK populations show altered receptor-defined subset balance in patients, consistent with dysregulated innate immune responses at the site of inflammation^27^ blood profiling further reports phenotypic and functional changes in NK cells, including shifts in activation state and gut-homing potential^28^. More recent clinical work also indicates broader NK-cell dysfunction in IBD, including dysregulated metabolic pathways that may compromise NK effector and regulatory functions^29^. Therefore, our finding that lower genetically regulated levels of NK CD56 bright cells are associated with higher Crohn’s disease risk supports the hypothesis that impaired innate immunoregulatory capacity, and consequent failure to maintain mucosal immune tolerance, contribute to Crohn’s pathogenesis.

## Discussion

In this study, we developed a framework to map the genetic regulation of immune cell-type proportions from genotyped single-cell RNA-seq data and to associate these cell types with autoimmune disease risk. Our main methodological innovation is a depth-weighted quasi-binomial GWAS that models proportions as bounded and overdispersed outcomes with depth-dependent precision. In the OneK1K dataset, this approach identified 47 genome-wide significant variants compared with 35 variants from a standard linear model. We further imputed genetically regulated proportions using polygenic scores and performed a summary-based cWAS across eight autoimmune diseases, yielding five significant cell type–disease associations (FDR < 0.1) and providing a direct bridge from genotype to disease without bulk deconvolution.

Specifically, our proportion GWAS identified a lead variant (rs4962627) located near *LHPP* that was associated with CD16^+^monocytes proportions. *LHPP* has been reported to be downregulated in colon tissue from patients with Crohn’s disease and ulcerative colitis, and in experimental colitis models, while *LHPP* loss alone is insufficient to induce or markedly worsen colitis, suggesting a modulatory or context-dependent role^30^. Together with our cWAS result that lower genetically predicted CD16^+^monocytes levels are associated with higher Crohn’s disease risk, these observations nominate *LHPP* as a candidate gene through which genetic control of monocyte composition may intersect with intestinal inflammation. These results motivate follow-up analyses to determine whether disease signals and cell-type signals share causal variants. Moreover, our framework is applicable to scRNA-seq samples with genotype data from other tissues or cellular compositions by integrating GRPs with GWAS summary statistics in cWAS analyses. In summary, we move beyond locus-disease catalogs to nominate disease-relevant immune cell types, offering a complementary view of disease biology and potential mechanisms underlying autoimmune susceptibility.

Our study has several limitations. First, although depth weighting improves robustness to sampling variability, cell-type proportion estimates can still be affected by technical factors such as cell capture biases and annotation uncertainty. These factors may introduce residual noise and attenuate effect estimates. Second, GRP-based cWAS analyses are constrained by the predictive performance of polygenic scores and by LD reference mismatches, and they do not, by themselves, establish that the same causal variants drive both composition and disease signals. In particular, significant GRP-disease associations can arise from correlated architectures, horizontal pleiotropy, or unmodeled confounding across linked variants, highlighting the need for further evaluation. Also, our analyses were restricted to genetically confirmed individuals of European ancestry, and extending the analyses to other populations would be valuable for assessing robustness and generalizability.

To strengthen mechanistic interpretation, future work could focus on using colocalization and fine-mapping to test whether cell-type proportions and disease GWAS signals at the same loci share causal variants, or on using Mendelian randomization to evaluate whether genetically regulated composition shifts plausibly lie on the causal path to disease. Beyond this, treating each cell-type proportion GWAS as a quantitative trait enables cell type-cell type genetic correlation analyses and module discovery to distinguish shared versus subset-specific genetic control. Cell type proportion GWAS summary statistics can also support polygenic predictors of GRPs that transfer beyond the discovery cohort, enabling phenome-scale translation and risk stratification based on genetically predisposed immune-composition profiles.

## Methods

### A weighted quasi-binomial GWAS model for cell type proportions

Traditional GWAS typically employs a linear model, regressing the phenotype on each SNP. However, it relies heavily on the Gaussian assumption, which may be violated by the right-skewed distributions of proportions for most cell types (Fig S1-S2). In addition, the proportions of cell types across individuals are highly variable, likely leading to overdispersion. To accommodate the excess variance typically observed in scRNA-seq-derived proportions, we introduce a dispersion parameter to improve flexibility and adopt a quasi-binomial regression model for cell type proportions:

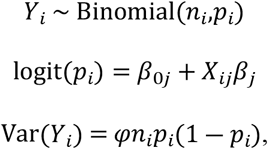

where *Y*_*i*_ represents the count of a specific cell type for individual *i, X*_*ij*_ is the genotype for individual *i* at SNP *j, β*_0*j*_ and *β*_*j*_ are the intercept and the genetic effect for SNP *j, n*_*i*_ denotes the total number of sampled cells, and *p*_*i*_ is the expected proportion of the given cell type. The additional dispersion parameter *φ* allows the variance to exceed the binomial assumption of *n*_*i*_*p*_*i*_ (1 − *p*_*i*_), thereby accounting for the overdispersion observed in cell-type proportions.

In scRNA-seq data analysis, sequencing depth across samples is crucial for the reliability of cell-type clustering. Greater sequencing depth may yield more precise estimates of cell-type proportions by reducing sampling variability. Thus, failure to implement appropriate weighting schemes may introduce bias in parameter estimation and lose statistical power. To address this concern, we adopt a weighting strategy based on the mean sequencing depth per cell per individual. Given that raw sequence depths varied widely, we linearly scaled each individual’s weights to mitigate the undue influence of samples with exceptionally high sequencing depth.

### Imputing GRP for association tests with complex diseases

To further identify the genetically regulated cell-type proportions associated with complex diseases, we followed the cWAS framework to impute GRPs for association studies^7^. By leveraging scRNA-seq samples with genotype data, our approach does not rely on bulk data or deconvolution methods used in the original cWAS framework. Instead, we used PRS to directly estimate the genetic weights of GRPs from GWAS summary statistics for cell-type proportions^31^. These pre-trained weights can be applied to external disease GWAS summary statistics for cWAS analysis. In brief, the cWAS summary statistics, *z*_*c, cwas*_, for cell type *c*, can be expressed as

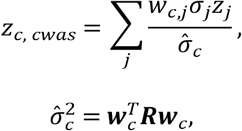

where *w*_*c,j*_ is the PRS weight of SNP *j* on cell type *c*, ***w***_*c*_ = (*w*_*c*,1_,…,*w*_*c,J*_)^*T*^ is a vector of PRS weights of all *J* SNPs, 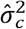 is the estimated variances of the GRP of the disease GWAS cohorts, 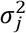 is the variance of the genotype of SNP *j, z*_*j*_ is the disease GWAS z score of SNP *j*, and ***R*** is the LD matrix.

Considering the huge computational cost to calculate the full LD matrix, we partition the SNPs into *B* approximately independent LD blocks, ***R*** ≈ *Diag*(***R***_(1)_, …, ***R***_(*B*)_), using LDetect^32^. Then, we can approximate the 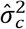 by

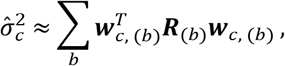

where ***w***_*c*, (*b*)_ is a vector of PRS weights of SNPs in the *b*th LD block. This approach reduces the computation complexity from *O*(*L*^*2*^) to approximately *O*(*L*^*2*^/*B*).

### OneK1K data processing

The OneK1K cohort is a single-cell RNA-seq resource of peripheral blood mononuclear cells (PBMCs) with matched donor-level genotypes and immune cell annotations^9^. We performed quality control on the genotype data, excluding variants with Hardy-Weinberg equilibrium (HWE) p-value ≤ 10^−6^ or minor allele frequency (MAF) ≤ 0.05. The original scRNA-seq data contained 31 distinct immune cell types. Our study focused on both fine and coarse cell types. To enhance the robustness of the analysis of genetic regulation in fine-labeled subtypes, we aggregated all dendritic cell subtypes, given their shared antigen-presenting functions, and excluded proliferative cell types and those with fewer than 2,000 total counts across all individuals. These approaches yielded 19 cell types with sufficient representation while minimizing potential noise from sparsely observed populations (Table S1; Fig S1). Furthermore, functionally related cell types were grouped into six coarse categories: B cells, CD4 T cells, CD8 T cells, Other T cells, Monocytes, and Natural Killer cells (Table S2; Fig S1). The proportion of each cell type was calculated as the number of cells belonging to that type relative to the total number of cells sequenced for each donor.

In the OneK1K data, some individuals exhibit abnormal proportions of certain cell types, potentially biasing the results. The disproportionate representation of specific cell types may be caused by several reasons: (1) Sampling or sequencing issues, like uneven cell recovery or biased cell sorting^33^; (2) Cells may have died or become stressed after sampling^34^; and (3) Cell types may be wrongly labeled. To remove such outliers, we fit a Beta distribution to the cell-type proportions across individuals for each cell type (Fig S2). A total of 17 donors with at least one cell type proportion exceeding the 99.9999th percentile of the fitted distribution were flagged as outliers and excluded from the study. This approach minimizes the influence of extreme proportions that may arise from technical artifacts or biological variability, thereby improving the robustness and interpretability of subsequent analyses. To mitigate the undue influence of samples with exceptionally high sequencing depth, we linearly scaled the weights for each individual from the original range of 1,806 to 5,969 reads per cell to a more flattened range of 0.9 to 1.1. Both models were adjusted for age, sex, and the first six principal components of genotypes.

### Cell-type Proportion PRS Calculation

GWAS summary statistics for cell type proportions were generated for both cell type-specific subsets and combined cell types. For each GWAS, we followed LD Hub guidelines^35^ for quality control by LD Score Regression (LDSC)^36^ to remove duplicate SNPs, strand-ambiguous SNPs (A/T and G/C), insertions and deletions (INDELs), and variants with effective sample sizes < 0.67 × the 90th percentile of the sample size distribution. We then applied PRS-CS-auto^31^ to infer posterior SNP effect sizes under a continuous shrinkage prior, using an LD reference panel from the 1000 Genomes Project with around 1.3 million HapMap3 SNPs in European populations. The resulting polygenic risk scores (PRS) were computed in the OneK1K dataset by summing the posterior effect sizes across all retained SNPs, weighted by the corresponding individual-level genotypes. When performing cWAS, we used LDetect^10^ to partition the genome into 2,197 LD blocks (∼1.6 centimorgan in width on average).

## Acknowledgments

Supported in part by NIH grants U01 HG013840 and U24HG012108.

## Supporting information captions

**Fig S1. Distribution of cell-type and cell-category proportions across individuals**.

**Fig S2. Beta-distribution fitting for outlier removal**.

**Fig S3. Manhattan plots for GWAS of coarse immune cell-type proportions**.

**Fig S4. Quantile–quantile (QQ) plots for generalized linear model association testing across fine-grained immune cell types**.

**Fig S5. Quantile–quantile (QQ) plots for linear model association testing across fine-grained immune cell types**.

**Fig S6. Heatmap of CWAS results across cell types and diseases. Table S1: Information for 19 cell types**.

**Table S2: Information for 6 integrated cell types**.

**Table S3: Genome-wide significant genetic variants (p<5×10^(−8)) identified by quasi-binomial model**.

## References

1 Dougherty JD, Schmidt EF, Nakajima M, Heintz N. Analytical approaches to RNA profiling data for the identification of genes enriched in specific cells. Nucleic acids research. 2010 Jul 1;38(13):4218–30.

2 Hekselman I, Yeger-Lotem E. Mechanisms of tissue and cell-type specificity in heritable traits and diseases. Nature Reviews Genetics. 2020 Mar;21(3):137–50.

3 Finucane HK, Reshef YA, Anttila V, Slowikowski K, Gusev A, Byrnes A, Gazal S, Loh PR, Lareau C, Shoresh N, Genovese G. Heritability enrichment of specifically expressed genes identifies disease-relevant tissues and cell types. Nature genetics. 2018 Apr;50(4):621–9.

4 Wang R, Lin DY, Jiang Y. EPIC: Inferring relevant cell types for complex traits by integrating genome-wide association studies and single-cell RNA sequencing. PLoS genetics. 2022 Jun 16;18(6):e1010251.

5 Jia P, Hu R, Yan F, Dai Y, Zhao Z. scGWAS: landscape of trait-cell type associations by integrating single-cell transcriptomics-wide and genome-wide association studies. Genome biology. 2022 Oct 17;23(1):220.

6 Yin M, Feng C, Yu Z, Zhang Y, Li Y, Wang X, Song C, Guo M, Li C. sc2GWAS: a comprehensive platform linking single cell and GWAS traits of human. Nucleic Acids Research. 2025 Jan 6;53(D1):D1151-61.

7 Liu W, Deng W, Chen M, Dong Z, Zhu B, Yu Z, Tang D, Sauler M, Lin C, Wain LV, Cho MH. A statistical framework to identify cell types whose genetically regulated proportions are associated with complex diseases. PLoS Genetics. 2023 Jul 31;19(7):e1010825.

8 Rumker L, Sakaue S, Reshef Y, Kang JB, Yazar S, Alquicira-Hernandez J, Valencia C, Lagattuta KA, Mah-Som A, Nathan A, Powell JE. Identifying genetic variants that influence the abundance of cell states in single-cell data. Nature genetics. 2024 Oct;56(10):2068–77.

9 Yazar S, Alquicira-Hernandez J, Wing K, Senabouth A, Gordon MG, Andersen S, Lu Q, Rowson A, Taylor TR, Clarke L, Maccora K. Single-cell eQTL mapping identifies cell type–specific genetic control of autoimmune disease. Science. 2022 Apr 8;376(6589):eabf3041.

10 Buniello A, Suveges D, Cruz-Castillo C, Llinares MB, Cornu H, Lopez I, Tsukanov K, Roldán-Romero JM, Mehta C, Fumis L, McNeill G. Open Targets Platform: facilitating therapeutic hypotheses building in drug discovery. Nucleic acids research. 2025 Jan 6;53(D1):D1467-75.

11 Bouillet P, Purton JF, Godfrey DI, Zhang LC, Coultas L, Puthalakath H, Pellegrini M, Cory S, Adams JM, Strasser A. BH3-only Bcl-2 family member Bim is required for apoptosis of autoreactive thymocytes. Nature. 2002 Feb 21;415(6874):922–6.

12 Chen MH, Raffield LM, Mousas A, Sakaue S, Huffman JE, Moscati A, Trivedi B, Jiang T, Akbari P, Vuckovic D, Bao EL. Trans-ethnic and ancestry-specific blood-cell genetics in 746,667 individuals from 5 global populations. Cell. 2020 Sep 3;182(5):1198–213.

13 Vuckovic D, Bao EL, Akbari P, Lareau CA, Mousas A, Jiang T, Chen MH, Raffield LM, Tardaguila M, Huffman JE, Ritchie SC. The Polygenic and Monogenic Basis of Blood Traits and Diseases. Cell. 2020;182(5):121411–3111.

14 Hindupur SK, Colombi M, Fuhs SR, Matter MS, Guri Y, Adam K, Cornu M, Piscuoglio S, Ng CK, Betz C, Liko D. The protein histidine phosphatase LHPP is a tumour suppressor. Nature. 2018 Mar 29;555(7698):678–82.

15 Barton AR, Sherman MA, Mukamel RE, Loh PR. Whole-exome imputation within UK Biobank powers rare coding variant association and fine-mapping analyses. Nature genetics. 2021 Aug;53(8):1260–9.

16 Rühlemann MC, Hermes BM, Bang C, Doms S, Moitinho-Silva L, Thingholm LB, Frost F, Degenhardt F, Wittig M, Kässens J, Weiss FU. Genome-wide association study in 8,956 German individuals identifies influence of ABO histo-blood groups on gut microbiome. Nature genetics. 2021 Feb;53(2):147–55.

17 Zaplana T, Miele S, Tolonen AC. Lachnospiraceae are emerging industrial biocatalysts and biotherapeutics. Frontiers in bioengineering and biotechnology. 2024 Jan 4;11:1324396.

18 Cantrell FL. Adverse effects of e-cigarette exposures. Journal of community health. 2014 Jun;39(3):614–6.

19 Karczewski KJ, Gupta R, Kanai M, Lu W, Tsuo K, Wang Y, Walters RK, Turley P, Callier S, Shah NN, Baya N. Pan-UK Biobank GWAS improves discovery, analysis of genetic architecture, and resolution into ancestry-enriched effects. MedRxiv. 2024 Mar 15:2024–03.

20 Bentham J, Morris DL, Cunninghame Graham DS, Pinder CL, Tombleson P, Behrens TW, Martín J, Fairfax BP, Knight JC, Chen L, Replogle J. Genetic association analyses implicate aberrant regulation of innate and adaptive immunity genes in the pathogenesis of systemic lupus erythematosus. Nature genetics. 2015 Dec;47(12):1457–64.

21 Okada Y, Wu D, Trynka G, Raj T, Terao C, Ikari K, Kochi Y, Ohmura K, Suzuki A, Yoshida S, Graham RR. Genetics of rheumatoid arthritis contributes to biology and drug discovery. Nature. 2014 Feb 20;506(7488):376–81.

22 International Multiple Sclerosis Genetics Consortium*†, ANZgene, IIBDGC, WTCCC2. Multiple sclerosis genomic map implicates peripheral immune cells and microglia in susceptibility. Science. 2019 Sep 27;365(6460):eaav7188.

23 De Lange KM, Moutsianas L, Lee JC, Lamb CA, Luo Y, Kennedy NA, Jostins L, Rice DL, Gutierrez-Achury J, Ji SG, Heap G. Genome-wide association study implicates immune activation of multiple integrin genes in inflammatory bowel disease. Nature genetics. 2017 Feb;49(2):256–61.

24 Forgetta V, Manousaki D, Istomine R, Ross S, Tessier MC, Marchand L, Li M, Qu HQ, Bradfield JP, Grant SF, Hakonarson H. Rare genetic variants of large effect influence risk of type 1 diabetes. Diabetes. 2020 Apr 1;69(4):784–95.

25 Koch S, Kucharzik T, Heidemann J, Nusrat A, Luegering A. Investigating the role of proinflammatory CD16+ monocytes in the pathogenesis of inflammatory bowel disease. Clinical & Experimental Immunology. 2010 Aug;161(2):332–41.

26 Cooper MA, Fehniger TA, Turner SC, Chen KS, Ghaheri BA, Ghayur T, Carson WE, Caligiuri MA. Human natural killer cells: a unique innate immunoregulatory role for the CD56bright subset. Blood, The Journal of the American Society of Hematology. 2001 May 15;97(10):3146–51.

27 Takayama T, Kamada N, Chinen H, Okamoto S, Kitazume MT, Chang J, Matuzaki Y, Suzuki S, Sugita A, Koganei K, Hisamatsu T. Imbalance of NKp44+ NKp46- and NKp44-NKp46+ natural killer cells in the intestinal mucosa of patients with Crohn’s disease. Gastroenterology. 2010 Sep 1;139(3):882–92.

28 Samarani S, Sagala P, Jantchou P, Grimard G, Faure C, Deslandres C, Amre DK, Ahmad A. Phenotypic and functional changes in peripheral blood natural killer cells in crohn disease patients. Mediators of Inflammation. 2020;2020(1):6401969.

29 Zaiatz Bittencourt V, Jones F, Tosetto M, Doherty GA, Ryan EJ. Dysregulation of metabolic pathways in circulating natural killer cells isolated from inflammatory bowel disease patients. Journal of Crohn’s and Colitis. 2021 Aug 1;15(8):1316–25.

30 Linder M, Liko D, Kancherla V, Piscuoglio S, Hall MN. Colitis is associated with loss of the histidine phosphatase LHPP and upregulation of histidine phosphorylation in intestinal epithelial cells. Biomedicines. 2023 Aug 1;11(8):2158.

31 Ge T, Chen CY, Ni Y, Feng YC, Smoller JW. Polygenic prediction via Bayesian regression and continuous shrinkage priors. Nature communications. 2019 Apr 16;10(1):1776.

32 Pickrell JK, Berisa T, Liu JZ, Ségurel L, Tung JY, Hinds DA. Detection and interpretation of shared genetic influences on 42 human traits. Nature genetics. 2016 Jul;48(7):709–17.

33 Yuan GC, Cai L, Elowitz M, Enver T, Fan G, Guo G, Irizarry R, Kharchenko P, Kim J, Orkin S, Quackenbush J. Challenges and emerging directions in single-cell analysis. Genome biology. 2017 May 8;18(1):84.

34 Arceneaux D, Chen Z, Simmons AJ, Heiser CN, Southard-Smith AN, Brenan MJ, Yang Y, Chen B, Xu Y, Choi E, Campbell JD. A contamination focused approach for optimizing the single-cell RNA-seq experiment. Iscience. 2023 Jul 21;26(7).

35 Zheng J, Erzurumluoglu AM, Elsworth BL, Kemp JP, Howe L, Haycock PC, Hemani G, Tansey K, Laurin C, Early Genetics and Lifecourse Epidemiology (EAGLE) Eczema Consortium, Pourcain BS. LD Hub: a centralized database and web interface to perform LD score regression that maximizes the potential of summary level GWAS data for SNP heritability and genetic correlation analysis. Bioinformatics. 2017 Jan 15;33(2):272–9.

36 Bulik-Sullivan BK, Loh PR, Finucane HK, Ripke S, Yang J, Patterson N, Daly MJ, Price AL, Neale BM. LD Score regression distinguishes confounding from polygenicity in genome-wide association studies. Nature genetics. 2015 Mar;47(3):291–5.

